# Hyperconcentrated Mucus Unifies Submucosal Gland and Superficial Airway Dysfunction in Cystic Fibrosis

**DOI:** 10.1101/2021.09.22.461306

**Authors:** Takafumi Kato, Giorgia Radicioni, Micah J. Papanikolas, Georgi V. Stoychev, Matthew R. Markovetz, Kazuhiro Aoki, Mindy Porterfield, Kenichi Okuda, Selene M. Barbosa Cardenas, Rodney C. Gilmore, Cameron B. Morrison, Camille Ehre, Kimberlie A. Burns, Kristen K. White, Tara A. Brennan, Henry P. Goodell, Holly Thacker, Henry T. Loznev, Lawrence J. Forsberg, Takahide Nagase, Michael Rubinstein, Scott H. Randell, Michael Tiemeyer, David B. Hill, Mehmet Kesimer, Wanda K. O’Neal, Stephen T. Ballard, Ronit Freeman, Brian Button, Richard C. Boucher

## Abstract

Cystic fibrosis (CF) is characterized by abnormal transepithelial ion transport. However, a description of CF lung disease pathophysiology unifying superficial epithelial and submucosal gland (SMG) dysfunctions has remained elusive. We hypothesized that biophysical abnormalities associated with CF mucus hyperconcentration provide a unifying mechanism. Studies of the anion secretion-inhibited pig airway CF model revealed elevated SMG mucus concentrations, osmotic pressures, and SMG mucus accumulation. Human airway studies revealed hyperconcentrated CF SMG mucus with raised osmotic pressures and cohesive forces predicted to limit SMG mucus secretion/release. Utilizing proline-rich protein 4 (PRR4) as a biomarker of SMG secretion, proteomics analyses of CF sputum revealed markedly lower PRR4 levels compared to healthy and bronchiectasis controls, consistent with a failure of CF SMGs to secrete mucus onto airway surfaces. Raised mucus osmotic/cohesive forces, reflecting mucus hyperconcentration, provide a unifying mechanism that describes disease-initiating mucus accumulation on airway surfaces and within SMGs of the CF lung.

## INTRODUCTION

Cystic fibrosis (CF) is characterized by abnormal transepithelial ion transport reflecting mutations in the cystic fibrosis transmembrane regulator (CFTR) gene (*1–3*). *In vitro* and *in vivo* studies of the superficial epithelia lining CF airway surfaces suggest dysfunctional CFTR-mediated ion transport produces hyperconcentrated mucus (*4–8*). Mucus hyperconcentration, defined as an increased mucus percent solids content (% solids) and/or mucin concentration, generates abnormal mucus biophysical properties, *e.g*., raised osmotic pressures and increased cohesive forces (*7, 9*). Osmotic compression of the periciliary layer (PCL) by the mucus layer in CF abolishes mucociliary transport, produces mucus accumulation on distal airway surfaces, and initiates the obstruction, infection, and inflammation that ultimately produce bronchiectasis (*8–11*).

However, there is controversy with respect to the contribution of CF submucosal gland (SMG) dysfunction to the proximal airway features of CF lung disease. Studies in CF pig tracheas described secretion of SMG mucus strands in response to maximal cholinergic stimulation (a mimic of cough) that are postulated to adhere to proximal CF airway surfaces, accumulate, and become a nidus for persistent bacterial infection (*12–17*). These data are juxtaposed to other studies that suggest CF SMG dysfunction reflects a failure to secrete SMG mucus with its robust antimicrobial components onto airway surfaces (*18–22*). Biophysical or clinical measurements that distinguish between these hypotheses have not been reported.

We hypothesized that normal SMG mucus has unique adaptations for host defense but, in common with superficial airway mucus, CF SMG mucus is hyperconcentrated, which limits secretion onto airway surfaces. Established methods were utilized for SMG mucus collection from freshly excised porcine and human airway tissues and multiple biochemical and biophysical properties of SMG mucus relevant to mucus transport measured. A biomarker selective for SMG secretion was identified and utilized as an index of SMG secretion *in vivo* in CF vs normal and disease control subjects.

## RESULTS

### SMG mucus from CF-mimicking pig airways exhibits raised concentrations and osmotic pressures

We first studied the osmotic pressures generated by normal vs CF-like SMG mucus from the anion transport inhibitor-treated wild-type (WT) pig model of CF (*23–26*). Because polymer biophysical properties are higher-order powers of polymer concentration, mucus concentration was measured in parallel (*5, 27*). Maximal acetylcholine stimulation, block of Cl^−^ secretion [bumetanide (Bum)], block of bicarbonate secretion [dimethylamiloride (DMA)], combined Bum/DMA block, or vehicle, in WT pig SMG mucus reproduced evidence of increased SMG mucus concentrations due to anion-transport inhibitor block of fluid secretion (% solids) (Fig.1A). Osmotic pressures were raised under conditions of fluid secretion block, paralleling % solids measurements (Figs.1B,C). Note, the anion transporter inhibitor protocols have opposite effects on SMG mucus HCO_3_^−^ concentrations, *i.e*., bumetanide raises HCO_3_^−^ concentrations, whereas bumetanide/DMA lowers HCO_3_^−^ concentrations, suggesting the increase in osmotic pressures generated by both inhibitor protocols reflected effects on mucus concentration and not HCO_3_^−^/pH (*28, 29*). Importantly, the osmotic pressure values measured in fluid block conditions are predicted to compress the PCL and slow/stop mucus transport (*5*).

**Figure 1.**
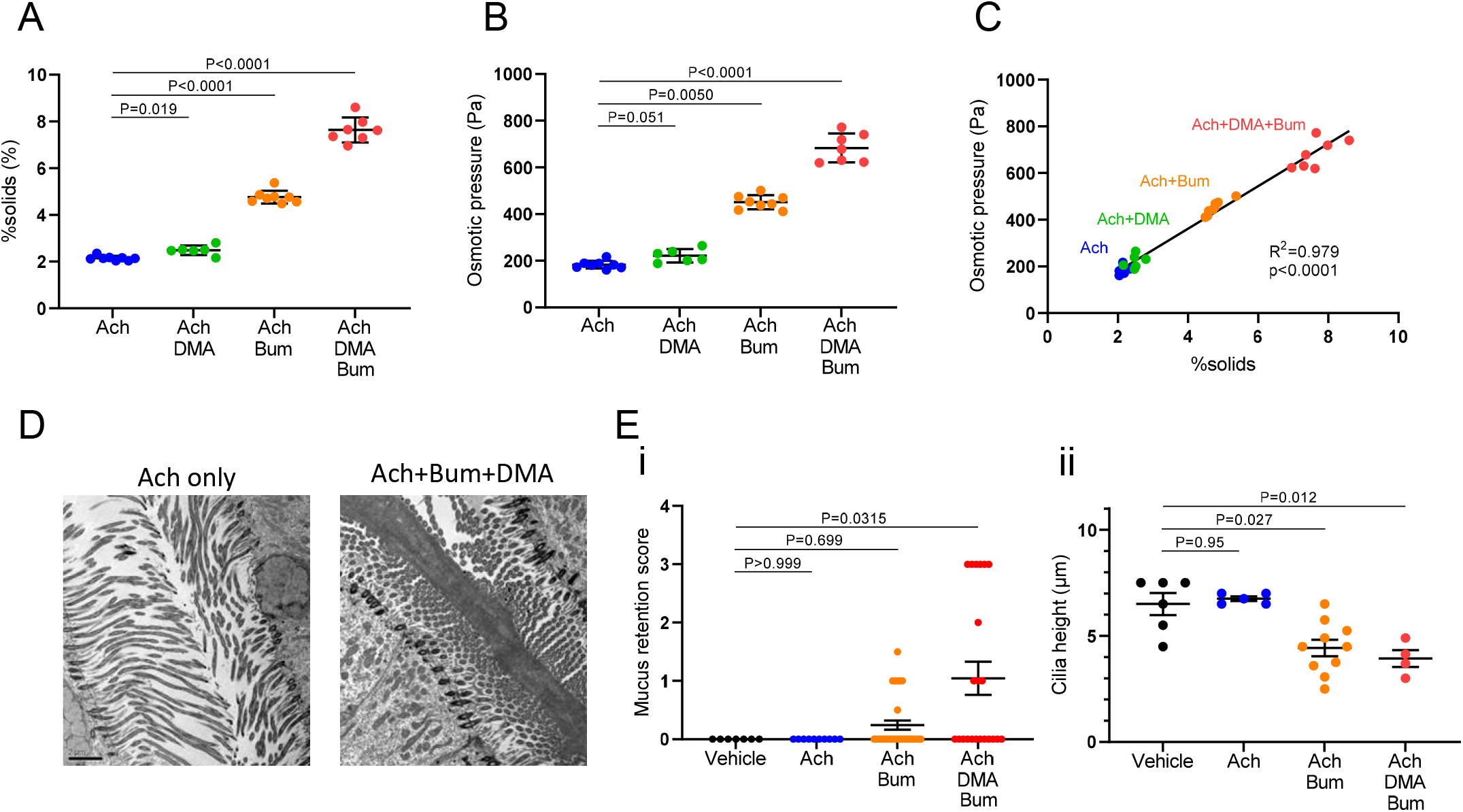
Studies of anion transport-inhibited porcine airways. **(A)** Solids concentration (% solids) of porcine submucosal gland (SMG) mucus after treatment with acetylcholine (Ach) without or with dimethylamiloride (DMA), bumetanide (Bum), or the combination of DMA and Bum. N = 6 to 8 per group. Welch’s ANOVA test followed by Dunnett’s multiple comparison test. **(B)** Osmotic pressure (in pascals, Pa) of porcine SMG mucus after treatment with Ach without or with DMA, Bum, or DMA/Bum. N = 6 to 8 per group. Welch’s ANOVA test followed by Dunnett’s multiple comparison test. **(C)** Linear correlation between %solids and osmotic pressure of pig SMG mucus. N = 29. **(D)** Transmission electron microscope (TEM) images of porcine SMG ducts after Ach treatment without or with DMA and Bum. **(E)** Quantitation of SMG (**i**) mucus retention and (**ii**) cilia height in ducts for each treatment condition. (**i**) N = 4 to 10 ducts from 3 airway specimens for each condition from 2 pigs. One to 8 regions of mucus retention per duct were evaluated. Each score was used for plot and statistical analyses. N = 72 regions from 25 ducts in total. Kruskal-Wallis test followed by Dunn’s multiple comparison. (**ii**) N = 4 to 10 ducts from 3 airway specimens for each condition from 2 pigs. One to 8 regions of cilia height per duct were measured. Mean of cilia height for each duct was used for plot and statistical analyses. Welch’s ANOVA test followed by Dunnett’s multiple comparison test.

### CF-mimicking maneuvers produce mucus retention and compressed cilia in SMG ducts

The presence of mucus and cilial height in SMG ducts of the control (Ach) and CF-mimicking (Ach+Bum+DMA) maneuvers were investigated using transmission electron microscopy (TEM). SMG mucus retention was observed in Bum/DMA (low gland fluid flow/anion secretion inhibition) conditions (Figs.1D,E). Cilia in glands treated with Bum/DMA were also compressed, suggesting ductal mucus retention reflected osmotic compression of cilia (Figs.1D,E) (*5*). Collectively, these data suggest that anion/fluid secretion inhibitors produced a hyperconcentrated pig SMG mucus that, via the osmotic compression of the ductal surface PCL, was retained in SMGs.

### Human CF SMG mucus exhibits increased concentrations, osmotic pressures, and cohesive strength

Consistent with the absence of CFTR anion/fluid secretory function in CF, CF SMG mucus was hyperconcentrated with increased osmotic pressures compared to non-CF SMG mucus (Figs.2A,B). Mucus concentrations correlated with osmotic pressure over ranges bracketed by CF and non-CF SMG mucus (Fig.2C). Studies designed to measure the forces required to “tear” mucus from gland orifices for transport up airway surfaces, *i.e*., cohesive forces (*30, 31*), identified elevated cohesive forces in CF compared to non-CF SMG mucus (Fig.2D).

**Figure 2.**
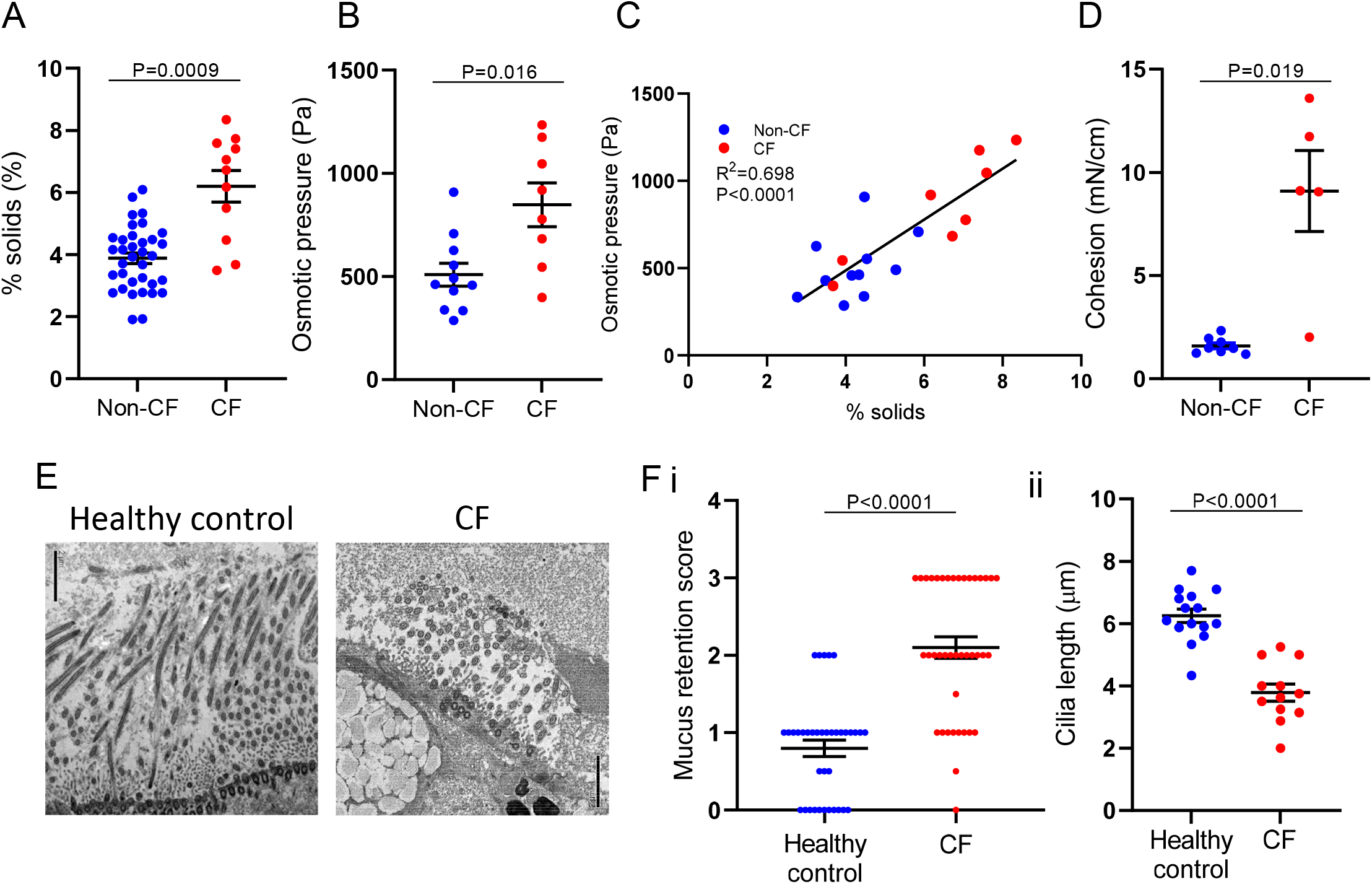
Studies of non-CF and CF human submucosal glands (SMGs) **(A)** Solids concentration (% solids) of human SMG mucus from non-CF (N = 30) and CF (N = 9) subjects. Welch’s t-test. **(B)** Osmotic pressure (Pa) of human SMG mucus from non-CF (N = 11) and CF (N = 9) subjects. Welch’s t-test. **(C)** Linear correlation between % solids and osmotic pressure of human SMG mucus. **(D)** Cohesive strength (mN/cm) of human SMG mucus from non-CF (N = 8) and CF (N = 5) subjects. Welch’s t-test. **(E)** SMG ducts of healthy control vs CF airways after acetylcholine treatment captured by transmission electron microscopy (TEM). Scale bar = 2μm. **(F)** Quantitation of human SMG (**i**) mucus ductal retention and (**ii**) cilia height in healthy control vs CF groups. (**i**) N = 37 regions from 15 healthy control ducts and N = 42 regions from 12 CF ducts. One to 8 regions of mucus retention per duct were evaluated depending on TEM image availability. Each score was used for plot and statistical analyses. Wilcoxon’s signed rank test. (**ii**) N = 15 ducts from 3 healthy control donors and N = 12 ducts from 3 CF donors. One to 7 regions of cilia height per duct were measured. Mean of cilia height for each duct was used for plot and statistical analyses. Welch’s t-test.

### CF SMGs exhibit mucus retention in ducts and compressed cilia

Consistent with the findings in CF-mimicking pigs (Figs.1D,E), human CF SMG ducts exhibited by TEM mucus retention and cilial compression (Figs.2E,F). These findings are consistent with previous reports of increased CF SMG mucus viscosity (*29*), and they predict that CF SMG mucus secretion is hindered by both osmotic compression of SMG mucus onto ductal surfaces and increased cohesive forces that limit mucus separation from ductal orifices.

### Insoluble strands as a unique feature of “normal” SMG mucus

We initiated studies of the properties of SMG mucus by comparing its composition with superficial airway epithelial mucus. Because SMG mucus in naïve porcine models has been reported to contain “strands” that maintain their structure after secretion onto airway surfaces, rather than dissolve into the ambient surface mucus layer (*12–15*), we investigated whether strand formation was unique to SMG mucus, *i.e*., not a feature of superficial epithelial mucus, and whether CF SMGs produced abnormal strands. Superficial HBE mucus dissolved fully into phosphate buffered saline (PBS), leaving behind no strand-like structures. In contrast, ~50% of Ach-stimulated non-CF human and pig SMG mucus failed to dissolve (Fig.3A). Addition of a non-ionic, non-denaturing detergent (IGEPAL) promoted dissolution of the insoluble SMG mucus component (Table.S1). Light scattering of the PBS-soluble SMG mucins demonstrated that SMG mucins were of similar size (Rg) as HBE mucins (Fig.3B). Light scattering of the insoluble SMG mucins was not possible. Instead, we immunostained the insoluble components of SMG mucus and identified SMG strands in mucus as comprised of MUC5B but not MUC5AC mucin (Fig.3C).

**Figure 3.**
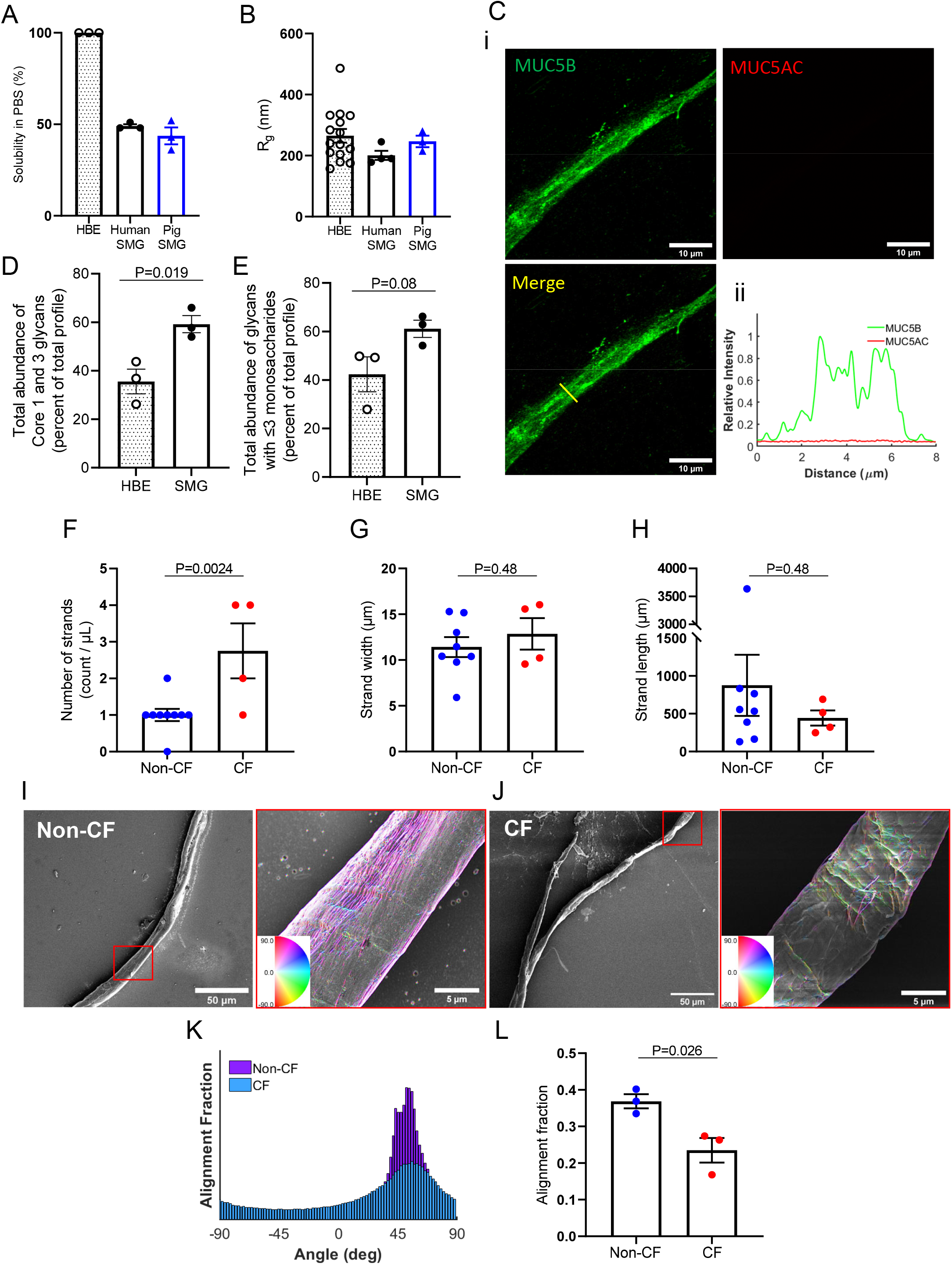
Submucosal gland mucus characteristics. **(A)** Solubility of human bronchial epithelial (HBE) cell culture mucus and human and porcine submucosal gland (SMG) mucus in phosphate buffered saline (PBS). **(B)** Radius of gyration (R_g_) of the soluble fraction of SMG mucus and HBE mucus determined by light-scattering. N = 10-15 for HBE, N=3 each for human and pig SMG mucus. Welch’s ANOVA test (P = 0.1003). **(C)**(**i**) Immunofluorescent staining of MUC5B and MUC5AC in non-CF SMG mucus strands. Scale bar = 10μm. (**ii**) Intensity of MUC5B and MUC5AC signals on the yellow line in the left panel. **(D)** Glycan abundance of less complex (Core 1 and 3) glycans in HBE vs SMG mucus. Unpaired t-test. N = 3 for each. (**E**) Abundance of glycans with shorter (≤ 3) monosaccharides in HBE vs SMG mucus. Unpaired t-test. N = 3 for each. **(F)** Relative number of strands found within SMG samples. Values were normalized by sample volume, imaged, and divided by the number of mean healthy strands. **(G)** Mean strand width of non-CF vs CF SMG mucus. **(H)** Mean strand length of non-CF vs CF SMG mucus. **(I)** Scanning electron microscopy (SEM) images of SMG strands from non-CF subjects. Inset falsely colored according to angular alignment of pixels quantified through OrientationJ. **(J)** SEM images of CF SMG strands. Inset falsely colored according to angular alignment of pixels. Samples from N = 9 (non-CF) and N = 4 (CF) donors. One to 5 strands per donor were measured. Mean of each donor was plotted. (**K,L**) Alignment of non-CF vs CF SMG mucus. **K** shows the histogram of relative alignment angles of SEM images in panels **I** and **J**. Non-CF sample shows increased alignment of surface features whereas CF strand does not. **L** shows the quantification of alignment fraction for each clinical sample based on decreased numbers of bins of panels **I** and **J** to account for slight deviations in angle from strand directionality. N = 3 each for non-CF and CF. (**F**) Mann-Whitney test. (**G**, **H**, and **L**) Unpaired t-test.

### Glycomics reveals SMG mucins contain shorter, less-branched glycans

Glycomics mass spectroscopy of soluble MUC5B from SMG vs superficial epithelial specimens from three matched donors was performed. SMG mucin-associated glycans contained a larger proportion of shorter, less-branched (Cores 1 and 3) carbohydrate structures than HBE mucins (Figs.3D,E,S1,Data.S1). Analyses of anionic O-glycans, in which the negative charge is imparted either as sulfate or sialic acid, were also performed. Sulfation of extended O-glycans, but not sialic acid, was reduced in SMG relative to HBE MUC5B (Figs.S1,S2,Data.S1,S2).

### CF SMG mucus strands exhibit distinct features

We visualized strands in the insoluble SMG component from non-CF and CF specimens using scanning electron microscopy (SEM). Consistent with CF SMG mucus hyperconcentration (Fig.2A), strand density was higher in CF SMG mucus (Fig.3F). The width and length of SMG strands were similar in non-CF and CF (Figs.3G,H). Alignment analyses demonstrated that non-CF SMG mucus strands contained extraordinarily long arrays of aligned bundles (Fig.3I), whereas CF SMG mucus strands exhibited randomly entangled bundles (Fig.3J). Quantitation of orientational order revealed less order, *i.e*., more heterogeneity, in CF strands (Figs.3K,L).

### Proline-rich protein 4 (PRR4) is a SMG-selective biomarker

Biomarkers selective for SMG secretions were investigated in human airway tissues. Four proteins, lactoferrin (LTF), zinc-α2-glycoprotein (AZGP1/ZAG), lysozyme (LYZ), and PRR4, were screened based on previous reports of selective SMG expression (*32–34*). RNA *in situ* hybridization (RNA-ISH) and immunohistochemistry (IHC) demonstrated that only PRR4 was selectively expressed in SMG serous cells and not in superficial epithelia lining large or small airways in healthy or diseased subjects (Figs.4A,B,S3). In contrast, LTF, AZGP1 and LYZ were variably detected in surface epithelia or infiltrating cells (Figs.S4A-F). qPCR data from freshly dissected specimens from different pulmonary regions confirmed the specificity of PRR4 for SMGs (Fig.S4G). Co-staining for PRR4 and MUC5B distinguished SMG serous and mucous acini, respectively (Fig.4C). SMG mucus passing through gland ducts exhibited co-positivity for PRR4 and MUC5B (Fig.4C, white arrow). In addition, SMG mucus strands exhibited co-localization of PRR4 and MUC5B (Fig.4D). Accordingly, PRR4 was selected as a biomarker to identify the contribution of SMGs to human airway secretions.

**Figure 4.**
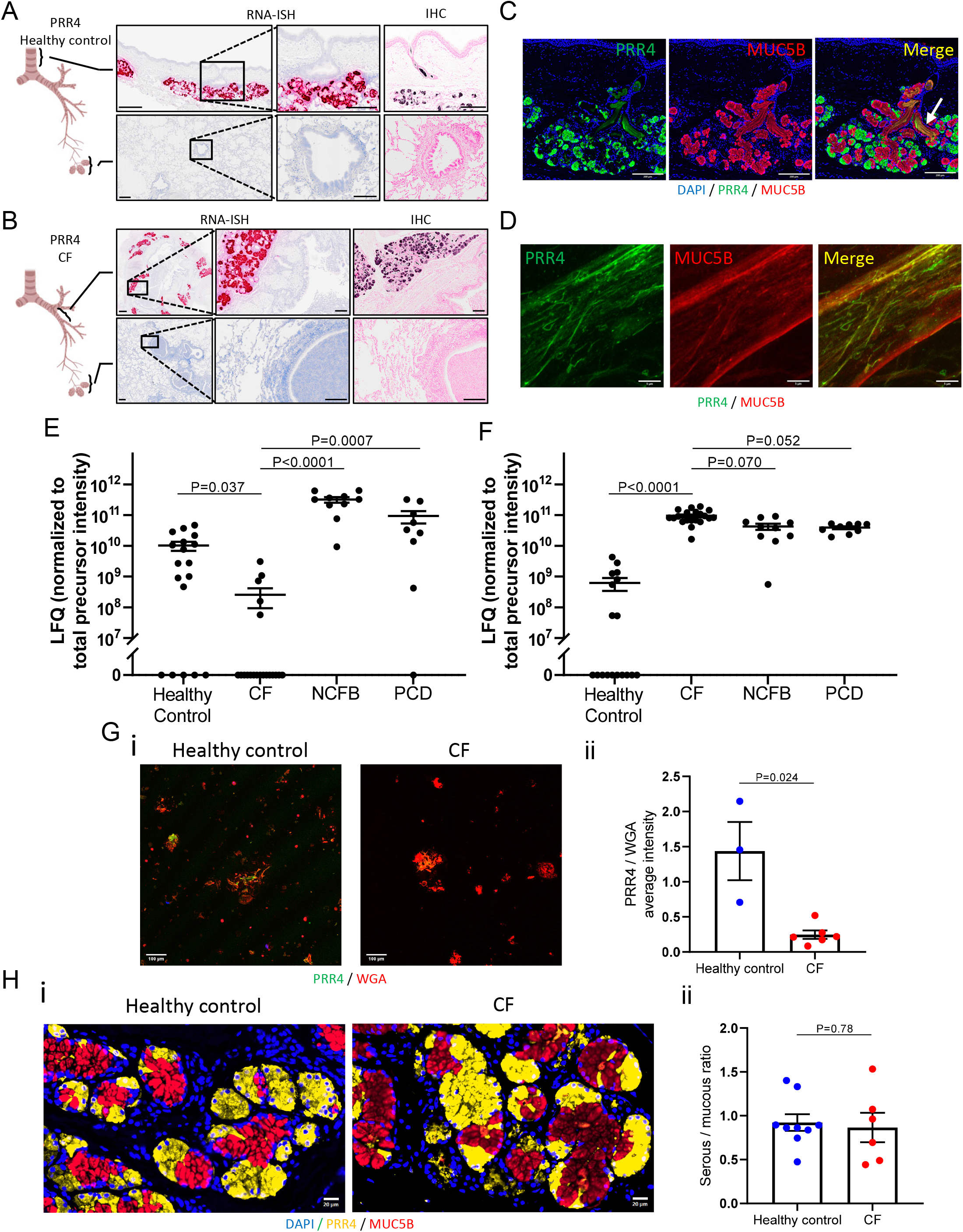
PRR4 in airways and clinical samples. **(A, B)** RNA *in situ* hybridization (RNA-ISH) and immunohistochemistry (IHC) for proline-rich protein 4 (PRR4) in (**A**) healthy control and (**B**) cystic fibrosis (CF) human airways. Bars: 500μm (low power images) and 200μm (magnified images). **(C)** Immunofluorescent co-staining of PRR4 and MUC5B in human proximal airways. SMG ductal mucus shows co-positivity for PRR4 and MUC5B (white arrow). Bars = 200μm. **(D)** Immunofluorescent co-staining of PRR4 and MUC5B in human SMG mucus strands. **(E,F)** Peptide intensities of (**E**) PRR4 and (**F**) human neutrophil elastase in induced sputum samples from healthy donors (N = 18), CF (N = 20), non-CF bronchiectasis (NCFB) (N = 10) and primary ciliary dyskinesia (PCD) (N = 9) measured by label-free quantitation (LFQ) utilizing mass-spectrometry. Kruskal-Wallis test followed by Dunn’s multiple comparison. **(G)** PRR4 content in BAL-obtained flakes. (**i**) Immunofluorescent co-staining of WGA and PRR4 for healthy control and CF mucus flakes (Bars = 100μm) and (**ii**) ratio of staining intensity of PRR4 to WGA in mucus flakes. N = 3 for healthy control, N = 6 for CF. Mann-Whitney test. (**H**) (**i**) Immunofluorescent staining of PRR4 and MUC5B for SMGs in healthy control vs CF tissues. Scale bar = 100μm. (**ii**) Quantification of serous cell (PRR4) vs mucous cell (MUC5B) signals in SMG in healthy control vs CF. N = 9 for healthy control, N = 6 for CF. Welch’s t-test.

### Sputum PRR4 level is reduced in CF but not in other diseases

Mass spectrometry (MS)-based proteomics analyses were performed on induced sputum samples collected from healthy controls and subjects with chronic airway diseases to investigate whether disease status affected the SMG contribution to airway secretions. CF sputum exhibited strikingly decreased PRR4 concentrations compared to both healthy (~ 30-fold reduction) and bronchiectatic disease controls (~ 300-fold reduction) (Fig.4E). This result did not reflect PRR4 degradation by human neutrophil elastase, as sputum HNE levels were similar in CF and NCFB/PCD subjects (Fig.4F) and PRR4 was not degraded by co-incubation with HNE *in vitro* (Fig.S5).

### Mucus flakes from CF subjects show reduced PRR4

Mucous flakes have been described in the insoluble material harvested by bronchoalveolar lavage fluid (BALF) from children with CF (*10*). Aliquots of these CF and disease control BALF samples were obtained and co-stained with a PRR4 antibody and wheat germ agglutinin (WGA) to localize mucin-associated lectins. The ratio of PRR4 to lectin staining was reduced in CF compared to healthy controls (Fig.4G).

### Healthy and CF airways have similar serous vs mucous cell proportion in SMG

To test whether reduced PRR4 levels in CF reflected reduced synthetic capacity, the proportion of serous vs mucous cells of SMGs in lungs resected from people with CF at time of transplant were quantified utilizing PRR4 and MUC5B immunostaining. Tissues from healthy and CF subjects exhibited similar ratios of PRR4-positive serous to MUC5B mucous cells (Fig.4H).

## DISCUSSION

CF airway surface epithelia exhibit hyperconcentrated mucus with raised osmotic pressures and cohesive strengths that are central to disease pathogenesis (*4, 5, 9*). However, less is known about native SMG mucus function and how it may be disturbed in CF. In this study, we tested the hypothesis that mucus hyperconcentration is a common defect driving CF pathophysiology in both airway surface epithelia and SMGs.

SMG mucus from pigs treated with anion/fluid secretion blockers was hyperconcentrated, consistent with previous reports by Ballard *et al* (*26, 35*) and Joo *et al* (*18, 19*) (Fig.1A). Human CF SMG mucus also was hyperconcentrated compared to non-CF SMG mucus (Fig.2A), consistent with defective fluid secretion *(16, 36)*. Importantly, elevated mucus osmotic pressures were measured in mucus from CF-mimicking pig and CF SMGs that paralleled increased mucus concentrations (Figs.1B,C,2B,C). Cohesive forces of human CF SMG mucus, also a function of increased mucus concentration, were also significantly elevated compared to control (Fig.2D). TEM investigations revealed that SMG ducts of CF-mimicking pig and human CF tissues exhibited both mucus retention and, as predicted from osmotic pressure measurements, cilial compression (Figs.1D,E,2E,F). These observations are consistent with previous reports of increased CF SMG mucus viscosity (*29*) and predict that CF SMG mucus secretion is hindered by both SMG mucus osmotic compression of ductal surfaces and increased cohesive forces that limit mucus separation from ductal orifices.

SMG mucus flows as a liquid but has been reported to contain “strands” that maintain their structure after secretion onto airway surfaces rather than dissolve into the ambient surface mucus layer (*14, 15*). We investigated the characteristics of SMG vs superficial epithelial soluble mucins and whether strand formation was unique to SMG mucus. Notably, superficial HBE mucus dissolved fully into PBS, leaving behind no strand-like structures, whereas ~50% of non-CF human and pig SMG mucus failed to dissolve into excess PBS (Fig.3A).

Glycomics analyses for the HBE vs SMG mucus from matched donors revealed differences in MUC5B glycosylation with potential implications for strand formation. First, SMG mucin glycans consisted of larger proportion of shorter, less-branched (Cores 1 and 3) carbohydrate structures than HBE mucins (Figs.3C,D,Data.S1). Mucin glycosylation with shorter glycans presents fewer hydroxyls for capture of water molecules through hydrogen bonding, a feature predicted to promote hydrophobic interactions between SMG MUC5B mucins and strand formation (37). Second, anionic O-glycans, in which the negative charge is imparted either as sulfate or sialic acid, were also measured. Sulfation of extended O-glycans, but not sialic acid, was reduced in SMG relative to HBE MUC5B, also consistent with a more hydrophobic, strand-prone SMG MUC5B mucin (Figs.S1,S2,Data.S1). The observation that a non-ionic, non-denaturing detergent (IGEPAL) promoted dissolution of the insoluble SMG strands is consistent with glycomics predictions (Table.S1). The co-localization of a unique SMG-secreted protein with SMG strands, *i.e*., PRR4 (discussed below), also suggests a role for protein cross-linkers in strand formation (Fig.4D).

Collectively, these data suggest specialized functions for surface vs SMG mucus in normal lung physiology. Superficial epithelial mucins likely dissolve rapidly upon secretion to replenish mucus cleared by transport. In contrast, strands secreted by SMGs maintain their structure on airway surfaces, *i.e*., are “permanent” gels (*10*). We speculate that strands aggregate with inhaled particles post cough-stimulated secretion to form collections (masses) of sufficient size to promote cough clearance in times of stress (*30*).

Distinct features of CF SMG mucus strands were identified (Fig.3). Consistent with CF SMG mucus hyperconcentration, strand density was higher in CF SMG mucus (Fig.3F). The width and length of SMG strands were similar in non-CF and CF mucus and strand width approximated the inner diameter of mucus tubules (~ 15μm diameter), suggesting that strand width is defined by distal SMG duct structures (Figs.3G,H) (*38*). Notably, SEM demonstrated that non-CF human SMG mucus strands contained extraordinarily long arrays of aligned bundles, whereas CF SMG mucus strands exhibited randomly entangled bundles (Figs.3I,J). Quantitation of orientational order revealed less order, *i.e*., more heterogeneity, in CF strands (Figs.3K,L). We speculate that decreased alignment of CF SMG strands reflects abnormal strand formation in a flow-limited, hyperconcentrated environment.

Finally, clinical investigations were performed to distinguish between the hypotheses that CF SMGs fail to secrete mucus vs secrete a strand-laden mucus that accumulates on airway surfaces (*17–19, 39*). PRR4 was identified as a biomarker selective for SMG secretion to identify and evaluate by mass spectroscopy the contribution of SMGs to airway secretions (Figs.4A-D,S4). NCFB and PCD subjects exhibited raised sputum PRR4 levels compared to healthy control subjects, consistent with SMG hypersecretion due to hypertrophied SMGs and/or increased stimulation of SMG secretion (*40*). In contrast, CF sputum exhibited strikingly decreased PRR4 concentrations compared to either normal subjects or bronchiectatic disease controls. Consistent with these results were data demonstrating that mucus flakes in CF BALF exhibited reduced PRR4 relative to mucin staining than control subjects. The reduced PRR4 levels in CF did not reflect PRR4 degradation by HNE (Fig.S5) or diminished PRR4 secretory capacity (Fig.4H). Collectively, these data suggest that the SMG contribution to CF pathogenesis reflects reduced CF SMG secretion onto, not accumulation of SMG mucus strands on, proximal airway surfaces.

Our study has several limitations. Because of the limited availability of fresh human airway tissues, specimens from non-CF diseased (*e.g*., smokers, chronic bronchitis) subjects were included in some studies (labeled as “Non-CF”). More detailed studies are necessary to fully characterize the SMG mucus properties in non-CF disease states to estimate the effect of tissue selection in this study. Another limitation is that maximal cholinergic stimulation of SMGs for 2 hours was required to generate sufficient SMG mucus volumes for analyses. Prolonged stimulation likely does not mimic the rapid (< 1 minute) stimulation of SMGs during cough. A final limitation is that we have not characterized the function of PRR4. Investigation of function of PRR4 and other proline-rich protein families (Fig.S4G) may reveal additional aspects of CF SMG pathogenesis.

In summary, SMG mucus exhibits unique characteristics configured for host defense, *e.g*., strand formation suitable for cough clearance. SMG mucus from human CF and CF-mimicking porcine airways is hyperconcentrated compared to non-CF SMG mucus. Hyperconcentrated SMG mucus exhibits increased osmotic pressures and cohesive forces predicted to retain mucus in SMG ducts and reduce secretion. The concentration of a SMG-selective secretory protein (PRR4) was lower in CF sputum than healthy controls, and strikingly lower than in subjects with non-CF bronchiectatic diseases, suggesting that CF SMG mucus secretion onto proximal CF airway surfaces is impaired. Accordingly, restoration of SMG host defense proteins/peptides on CF airway surfaces early in life may be therapeutic (*41*). Finally, these data suggest that increased mucus osmotic/cohesive forces, a product of abnormal CFTR-dependent mucus hydration (hyperconcentration), provide a unifying mechanism to describe the muco-obstructive distal and proximal airway components of CF lung disease.

## MATERIALS AND METHODS

### Human airway dissection

Human tissues were provided by the University of North Carolina (UNC) Tissue and Cell Culture Procurement Core under protocols approved by UNC Office of Research Ethics Biomedical Institutional Review Board (IRB numbers 03-1396 and 17-3281). Non-CF airway tissues were obtained from lungs of donors that could not be used for transplant. CF airways were obtained from lungs resected from CF patients undergoing transplantation at UNC Hospitals. Excised tissues were transported in saline on ice, and all studies were conducted within 12 to 36 hours of lung removal. Proximal airways (2^nd^ to 4^th^ generation bronchi) were manually dissected. Intraluminal mucus was removed by washing with Ringer’s bicarbonate solution until the flushed solution appeared clear, and residual fluid was collected as completely as possible with positive-pressure pipettes.

### Pig airway dissection

All procedures with swine performed at the University of South Alabama were conducted under protocols approved by the University of South Alabama Institutional Animal Care and Use Committee in compliance to the US Public Health Service *Policy on Humane Care and Use of Laboratory Animals*. Pig bronchial dissection was performed as previously described (*26*). For the collection of SMG mucus for solubility and light scattering assays performed at UNC, freshly excised porcine airway tissues were obtained from a local slaughterhouse. Main stem or segmental bronchi were dissected in the lab.

### Gland mucus collection

Pig and human airway SMG mucus collection was performed using established protocols with minor modifications (*26, 36*). The proximal end of an isolated airway was cannulated with polyethylene tubing and ligated with sutures. All distal branches of the excised preparation were tightly ligated with suture material to physically separate the airway lumen from the incubation buffer, enabling SMG mucus to be secreted into the lumen without contamination by incubation buffer (Krebs-Ringer bicarbonate, pH 7.4, under continuous gassing with 95% O_2_ and 5% CO_2_ at 37°C). For the porcine main stem to segmental bronchi, both proximal and distal edges were sealed with silicon plugs. Acetylcholine chloride (A625, Sigma) was added to the incubation solution at a concentration of 10μM to stimulate SMG secretion. For porcine experiments, dimethylamiloride (A4562, Sigma) and/or bumetanide (B3023, Sigma) were used together to block bicarbonate and/or chloride secretion, respectively. After 2-hour incubation interval with acetylcholine with or without anion transport inhibitors, the bronchial tissues were removed from the buffer solution and decannulated. After wiping the outer surface of the preparation to remove buffer solution, the airway was opened longitudinally. Mucus secreted into the lumen was collected by aspiration with a 10μl positive pressure pipette. In each experiment, % solids concentration was measured immediately (described below), and the remaining sample was kept frozen at −80°C until further investigation.

### % solids measurement

Percent solids concentrations (% solids) of human and porcine SMG mucus (3 to 5 μl) were measured gravimetrically as described previously (*42*). The mucus aliquots were weighed on a pre-weighed piece of aluminum foil. The samples were heated at 80□ in an oven overnight to allow the liquid content to evaporate completely. The final weight of the dried foil and mucus sample was determined and solid weight% (% solids) was calculated.

### Osmotic pressure measurement

Osmotic pressures of human and pig SMG mucus (20-30 ul) were measured with a custom-designed osmometer incorporating a 10-kDa membrane as previously reported (*5*). Twenty to 30 μl of SMG mucus was placed into the fluid chamber, and the steady-state osmotic pressure of mucus was acquired by the pressure transducer placed on the other side of the osmotic membrane.

### Cohesion measurements

The cohesive strength of human SMG mucus samples (20-40 ul) was measured using a custom-designed peel test device (*30, 31*). Twenty to 40 μl of a SMG mucus sample was placed on a mucus-binding mesh and the peeling-force to tear the mucus apart was measured.

### TEM for porcine and human airway tissues

Airway tissues were immersed in 2% paraformaldehyde/2.5% glutaraldehyde/0.15M sodium phosphate, pH 7.4 and stored for several days at 4□. After post-fixation for 1 hour with buffered potassium ferrocyanide-reduced osmium (1% osmium tetroxide/1.25% potassium ferrocyanide/0.15M sodium phosphate) the samples were washed in deionized water, then dehydrated through a graded ethanol and propylene oxide (*43*). The samples were infiltrated with a 1:1, then 1:2 mixture of propylene oxide and Polybed 812 epoxy resin for 4 hours and infiltrated overnight in 100% resin (08792-1, Polysciences). Samples were transferred to embedding molds and polymerized at 60□ overnight. Using a diamond knife, 1μm semi-thin sections were cut, mounted on slides, and stained with 1% toluidine blue to examine by light microscopy and isolate regions of interest. Ultrathin sections (70-80 nm) were cut with a diamond knife and mounted on 200 mesh copper grids followed by staining with 4% aqueous uranyl acetate for 12 minutes and Reynold’s lead citrate for 8 minutes (*44*). The samples were observed using a JEOL JEM-1230 transmission electron microscope operating at 80kV (JEOL USA) and images were acquired with a Gatan Orius SC1000 CCD Digital Camera and Gatan Microscopy Suite 3.0 software (Gatan). Quantitative measurement of cilial height and semi-quantitative scoring of mucus retention in SMG ducts were performed by a blinded observer. Cilial height from the ductal cell surface to the tip of the cilia was quantified. Mucus retention was scored from 0 to 3 with 0.5 intervals.

### SEM of SMG mucus

SMG mucus was diluted 10-fold in PBS and mixed for two hours at room temperature (RT). Samples were allowed to sediment onto an APTES-coated surface for 15 minutes before being fixed with 2% paraformaldehyde, 2.5% glutaraldehyde in PBS for another 15 minutes. The samples were washed three times with sterile water and dehydrated into ethanol using increments of 10% in ten-minute increments. The samples were then critically point dried and mounted onto a SEM specimen holder using copper tape, coated with 5nm AuPd (108 Auto, Cressington) and imaged on SEM (FEI Helios Nanolab 600) using 5kV accelerating voltage. Analysis of orientation was performed using OrientationJ (http://bigwww.epfl.ch/demo/orientation/) using a method adapted from Nerger *et al* (*45*).

### Human airway surface mucus production

HBE cells were isolated from non-smoker donors without respiratory diseases by the UNC Tissue and Cell Culture Procurement Core as described (*46*). For glycomics comparison of SMG vs HBE mucus, isolated HBE primary (passage 0) cells, donor-matched to tissue sections from which SMG mucus was collected, were seeded on collagen (5005, Advanced BioMatrix)-coated Transwell inserts (3460, Corning) at a density of 0.3 million cells per insert. For solubility and light scattering studies, HBE primary cells were expanded as previously described (*47*). Passage 1 or 2 cells were seeded on collagen-coated Transwell inserts at a density of 0.3 million cells per insert. The cells were cultured under submerged condition until confluence followed by air-liquid interface (ALI) culture with UNC ALI media (*48*). After a full differentiation confirmed by ciliation under light microscopic observation, secreted mucus was allowed to accumulate on apical surface for up to three weeks and collected by washing with PBS. The HBE mucus was pooled from multiple codes and then concentrated with ultrafiltration using 50K cutoff filter (UFC905096, Millipore) to the %solids of 4%.

### Solubility measurements

Collected SMG and HBE mucus was diluted 40-fold with PBS and kept at 4°C for 4 days to allow the mucus to solubilize. After the 4-days, HBE mucus was completely dissolved, whereas the SMG mucus dissolved only partly in the PBS. Undissolved portions were diluted in PBS and %solids of the insoluble part was calculated. The SMG supernatant, containing the soluble component of SMG mucus, was collected for light scattering. The PBS-insoluble SMG mucus was exposed to surfactant (1 mM in PBS, IGEPAL CA-630, Sigma) and solubility was reexamined 48 hours later.

### Light scattering

Static light scattering for soluble components of SMG and HBE mucus were performed on a research goniometer (BI-200SM, Brookhaven Instruments), equipped with a 640nm laser. A series of solutions with different concentrations were measured at scattering angles between 40° and 150°. The data were plotted as a Zimm plot, from which the radius of gyration (R_g_) of the solutes were extracted. The lognormal size distribution of the solutes was obtained by cumulant analysis of the correlation function between 100 nsec and 10 msec. Prior to measurements, all solutions were filtered through a 0.45 μm hydrophilic PVDF syringe filter (Titan 3, Thermo Scientific) to remove dust particles and large cell debris.

### Analysis of mucin O-glycosylation

O-glycans from HBE and SMG mucus were released by reductive β-elimination and analyzed by nano-spray ionization multi-dimensional mass spectrometry (NSI-MSn) as described (*49, 50*). Briefly, mucus samples were precipitated from cold 80% aqueous acetone, dried, and resuspended for reductive β-elimination. Following release and clean-up, O-glycans were permethylated and recovered by water-dichloromethane (DCM) extraction. Non-sulfated and sulfated O-glycans were recovered in the DCM and aqueous phases, respectively. The two phases were separated, dried, and resulting glycans were analyzed by NSI-MSn. MS data was acquired in both negative and positive ion modes to detect sulfated and non-sulfated glycans, respectively. For quantification of permethylated, non-sulfated glycans, a known amount of maltotetraose permethylated with isotopically heavy methyliodide was added to each O-glycan sample, and mucin glycan amounts were measured in reference to the standard (*51*). For sulfated glycans, the same quantification was not possible due to ionization suppression caused by the sulfate moiety. Therefore, sulfated glycans were quantified based on the signal intensity measured in the mass spectrometer. Graphical representations of monosaccharide residues are presented in accordance with the broadly accepted Symbolic Nomenclature For Glycans (SNFG) and O-glycan analysis was performed according to the MIRAGE guidelines (*52*).

### Tissue slide preparation for RNA-ISH and IHC

Human airway and lung tissues were provided by the UNC Tissue and Cell Culture Procurement Core. The donors included healthy controls (non-smokers without any history of respiratory diseases), CF, asthma, NCFB and PCD. Tissues were collected from multiple generations, *i.e*., trachea, main bronchi, and distal lungs with bronchioles. The dissected tissues were fixed with 10% neutral buffered formalin for 24-36 hours and embedded in paraffin. The formalin-fixed paraffin-embedded (FFPE) blocks were cut to 5 μm.

### RNA-ISH

RNA-ISH was performed using the RNAscope^®^ 2.5 HD Reagent Kit-RED (322350, Advanced Cell Diagnostics) according to the manufacturer’s instruction and published protocols (*47*). Probes targeting *PRR4* (522791), *LTF* (425101), *AZGP1* (401861) and *LYZ* (421441), in addition to positive and negative controls (310041 and 310043, respectively), were hybridized at 40□ for 2 hours in the HybEZ oven (241000) followed by signal amplification and washing. Signals were visualized by Fast Red followed by counterstaining with hematoxylin. The probe targeting human housekeeping gene Ubiquitin C (*UBC*, 310041) served as a positive control to evaluate RNA quality. A bacterial gene, Bacillus subtilis dihydrodipicolinate reductase (*DapB*, 310043) served as negative control. The images were acquired using an Olympus VS200 slide scanner light microscope with a 60x 1.42 N.A. objective.

### Immunostaining

FFPE tissue sections were baked at 60°C for 2□hours, deparaffinized with xylene for 10 minutes three times, and rehydrated with graded ethanol. Antigen retrieval was performed by boiling slides in 0.1□M sodium citrate pH 6.0 (1200W microwave settings: 100% power for 6.5□minutes, followed by 60% power for 6□minutes twice). After cooling, quenching of endogenous peroxidase was performed with 0.5% hydrogen peroxide in methanol for 15□minutes. Then slides were blocked with Blocking One (03953-95, Nacalai Tesque) at RT for 30 minutes. The primary antibodies rabbit anti-PRR4 (PA5-59883, Invitrogen, 1:2500), anti-LTF (10933-1-AP, Proteintech, 1:1000), anti-ZAG (13399-1-AP, Proteintech ,1:1000) and anti-LYZ (MA5-32154, Invitrogen, 1:1000) were applied to slides and incubated at 4□ overnight. Sections were washed with PBS and incubated with biotinylated donkey anti-rabbit IgG secondary antibodies (711-065-152, Jackson ImmunoResearch, 1:400) at RT for 1 hour. Slides were treated with Vectastain Kit (PK-4000, Vector Laboratories) according to manufacturer’s instruction. Sections were stained with 3,3’-diamino benzidine and counterstained with Fast Red. Images were acquired as described above. For immunofluorescent co-staining, tissue sections were treated with rabbit anti-PRR4 antibody and mouse anti-human MUC5B antiserum (raised in-house, M-25, 1:1000) at 4□ overnight followed by fluorescent staining with Alexa Fluor Plus 647-conjugated donkey anti-rabbit IgG (A32795, Invitrogen, 1:1000) and Alexa Fluor® Plus 555-conjugated donkey anti-mouse antibody (A32773, Invitrogen, 1:1000) at RT for 30 minutes. Arter autofluorescence quenching (SP-8400-15, Vector Laboratories), tissue was mounted with DAPI-containing medium (P36931, Invitrogen). Fluorescent images were acquired using Olympus VS200 scanner with DAPI (488 nm), Cy3 (555 nm) and Cy5 (647 nm) filters. Isotype controls were prepared using rabbit and/or mouse IgG (011-000-003 and 015-000-003, Jackson ImmunoResearch). Volume ratio of serous (PRR4 positive) cells to mucous (MUC5B positive) cells volume in SMGs was quantified with Olyvia (Olympus) and VisioPharm (VisioPharm) software.

### Immunocytochemistry of SMG mucus

SMG mucus was diluted 10-fold in PBS and mixed for 2 hours at RT. Samples were allowed to sediment onto an APTES-coated surface for 15 minutes before being fixed with 2% PFA for another 15 minutes. Samples were washed 3 times with PBS. Samples were then incubated with PRR4 antibody and MUC5B antisera or MUC5B antibody (H-300, Santa Cruz, 1:1000) and MUC5AC antibody (45M1, Invitrogen, 1:1000) at 4□ overnight followed by washing 3 times with PBS and incubated with secondary antibodies overnight at 4□. Samples were washed again and stained with DAPI (D9542-5MG, Sigma, 1:10,000) for 15 minutes, and then washed and mounted onto slides (9990402, Fisher Scientific). Images were acquired with a Zeiss 880 confocal microscope. To view co-localization of MUC5B and MUC5AC, Airyscan acquisition mode was used for imaging (Fig.3C).

### RNA isolation from multiple regions of human airways

Freshly excised airway tissues from healthy non-smoker donors were dissected under light microscopy. Small (~ 1 cm^2^) sections that contained surface and submucosal tissue were dissected from trachea and main bronchus. Small bronchioles were carefully dissected from surrounding alveolar tissue. Distal lung parenchyma sections of 0.5 to 1cm^2^ in size were cut as alveolar tissues. Dissected specimens were minced with a homogenizer, and total RNA extracted with TRI reagent (T9424, Sigma) and an RNA extraction kit (R2052, Zymo Research).

### cDNA synthesis and RT-qPCR

200ng of RNA was used for synthesizing cDNA (1708840, Bio-Rad) according to the manufacturer’s instructions. qPCR was performed with QuantStadio 6 (Thermo Fisher). Taqman PCR probes (Life Technologies) used for the investigation of SMG markers included *PRR4* (Hs00200615_m1), *PRB1* (Hs00818764_m1), *PRB3* (Hs00818925_m1), *PRB4* (Hs00864002_m1), *PRH2* (Hs00818765-mH), *LTF* (Hs00914334_m1), *AZGP1* (Hs00426651_m1) and *LYZ* (Hs00426232_m1). A marker for distal bronchioles and alveoli, *SFTPB* (Hs00167036_m1) was measured to assure airway regions were obtained from. *TBP* (Hs00427620_m1) was chosen for the internal control. All probes were designed to span exons.

### Label-free quantification mass spectrometry for induced sputum

Induced sputum samples were obtained from the UNC sample repositories (IRB numbers 16-3142 (NCFB), 15-2431 (healthy controls and CF) and 02-0948 (PCD)). One hundred μl of samples were prepared with the Filter Aided Sample Preparation (FASP) method. Briefly, samples were denatured with 800 μl of 6M GuHCl (pH 8.0) and reduced by adding DTT to a final concentration of 20 mM for 1 hour at 65□. After reduction, samples were alkylated with iodoacetamide (I1149, Sigma) at a final concentration of 50 mM for 1 hour at 25□ in the dark. Samples were centrifuged at 14,000 g for 10 min, and the filter was washed twice with50 mM ammonium hydrogen carbonate (NH_4_HCO_3_). 0.5 μg modified trypsin (proteomics grade, T6567, Sigma) was added and samples were incubated for 18 hours at 37□. The peptides solutions were concentrated by vacuum centrifugation and dissolved in 30 μL 0.1 % formic acid. Mass spectrometry was performed with a Dionex ultimate 3000 RSLCnano system coupled to a hybrid quadrupole orbitrap mass spectrometer with a Nano spray source (Thermo Scientific Q-Exactive). Samples (3 μL) were loaded into a trap column (Acclaim PepMap 100, 100 μm x 2 cm, nanoViper C18 5 μm 100 Å), at 5 μL/min with aqueous solution containing 0.1 % (v/v) trifluoroacetic acid and 2 % acetonitrile. The column used for peptides separation was an Acclaim PepMap RSLC, 75 μm x 15 cm, nanoViper C18 2 μm 100 Å. LC – MS runs were 120 minutes long, with two liquid phases: 97% water 3% Acetonitrile 0.1% Formic Acid (buffer A) and Acetonitrile 0.1% Formic Acid (buffer B). The gradients (% of buffer B) of the run (120 minutes) applied were; 0 to 7 minutes: 4%; 7 to 90 minutes: 30%; 90 to 105 minutes: 80%; and 105 to 120 minutes: 4%.

Data were acquired at a resolution of 70,000 at m/z 200, target AGC value of 5e5, maximum fill times of 200 ms, a multiplex degree of 6 with an isolation width of 3 m/z.

Proteins were identified by searching against the most current human database with Proteome Discover 1.4, setting a maximum of two missed cleavage sites for trypsin, oxidation of methionine (+15.995 Da) as a dynamic modification, and carbamidomethylation of cysteine (+57.0210 Da) as static a modification. Proteins were quantified using Scaffold (Proteome Software), using the normalized total precursor intensity, with 95% probability by the Scaffold Local FDR algorithm.

### PRR4 and WGA staining for airway mucus flakes

Mucus flakes were obtained from BALF from healthy non-smoker adult subjects and school-age CF subjects as previouswily reported under IRB-approved protocol at UNC Chapel Hill (IRB numbers 91-0679 and 11-1445) (*53*). All studies were approved by the UNC Ethics Committee and informed consents from the subjects or parents obtained prior to any study procedures. BALF (10 μl) was applied to glass microscopy slides using a cytocentrifuge (StatSpin CytoFuge 2, Beckman Coulter). Slides were fixed with 10% neutral buffered formalin, washed with PBS, and blocked with 3% bovine serum albumin at RT for 1 hour. PRR4 and mucus were immunohistochemically stained by PRR4 antibody (1:1000) and biotinylated WGA (BK-1000, Vector Laboratories, 1:1000), respectively, at 4°C overnight. Slides were washed with PBS three times, followed by incubation with secondary antibodies (Alexa Fluor® Plus 488-conjugated donkey anti-rabbit IgG (A32795, Invitrogen, 1:1000) and 594-conjugated streptavidin (U0291, Abnova, 1:1000)) and DAPI (D3571, Invitrogen, 1:1000) at RT for 1 hour. The slides were then washed with PBS and mounted with Fluorsave mounting media (345789, Millipore). The staining images were obtained with an Olympus FV1000 confocal microscope using a 20x objective and uniform settings. Average PRR4 intensity was normalized to overall mucus signal by dividing by the average WGA intensity.

### Production of recombinant PRR4 protein

PRR4 cDNA was obtained from ORF clone plasmid (OHu15631, GenScript). PRR4 cDNA without a signal peptide sequence (corresponding to amino acids 17 to 134) was cloned into the pM-secSUMOstar Vector (7121, LifeSensors). SUMOstar-PRR4 vectors were transfected into Expi293 cells (1 mg of DNA per liter of transfection) per manufacturer instructions, with culture supernatants harvested and sterile-filtered on day 3 post-transfection. Supernatants were then concentrated and buffer-exchanged by tangential-flow filtration into TALON-binding buffer (50 mM sodium phosphate pH 7.4, 500 mM NaCl, 5 mM imidazole, 0.05% sodium azide, 10% glycerol), then loaded onto hand-packed 1 ml TALON (635507, Takara Bio) columns recirculating overnight at 1.5 ml/min. Twenty-four hours later, columns were changed to flow-through, and run until all supernatant was loaded. Columns were then washed with 20 column volumes binding buffer, followed by binding buffer with 10 mM imidazole. Proteins were then eluted with 20 CV binding buffer supplemented with 150 mM imidazole. Fractions were assessed by SDS-PAGE, and those containing bands corresponding to His-SUMO-tagged PRR4 (10 mM wash and elution) were pooled. Pooled fractions were then dialyzed against PBS with 2 mM DTT (3.5 kDa MWCO). SUMOstar protease (SP4110, LifeSensors; 10 units/mg substrate) was added to dialyzed protein and the reaction was incubated overnight at 4°C to cleave the His-SUMO tags. Cleaved proteins were then passed back over the 1 ml TALON columns to separate the His-SUMO tags from the cleaved PRR4 proteins. Subtractive columns were washed and eluted as described above. Fractions were again assessed by SDS-PAGE to confirm separation, with flow-through and 5 mM imidazole fractions pooled, concentrated, and dialyzed into PBS (3.5 kDa MWCO). The final purified protein was confirmed to be PRR4 by both western blotting and mass spectrometry.

### Confirmation of no degradative effect for PRR4 by neutrophil elastase

To quantitate the potential effects of proteolysis on the proteomics results, endotracheal tube (ETT) mucus samples were treated with designated quantities of HNE (10-50 ug/ml, 324681, Sigma). ETT mucus was obtained as previously reported from endotracheal tube tips following post-surgical extubation at UNC General Hospital under UNC IRB approved protocols (IRB number 17-2211) (*54*). Aliquots of ETT mucus (500uL) were treated with 10, 20 and 50 ug of HNE and rotated/mixed for 30 minutes, and then incubated at 37□ for overnight. PBS was added to the control. PRR4, MUC5B and MUC5AC peptide intensities were measured by mass spectrometry as described above. (Fig.S5).

### Validation of mouse anti-MUC5B antiserum for immunostaining

The mouse anti-MUC5B antiserum raised in-house was not previously published. As a validation, we performed immunohistochemistry of MUC5B for human tracheal serial sections with the antiserum (M-25, 1:1000) and previously established commercial rabbit anti-MUC5B antibody (H-300, Santa Cruz, 1:1000). Staining with these antibody and antiserum appeared identical, confirming the sensitivity and specificity of the novel antiserum (Fig.S6).

### Statistical analyses

Statistical analyses and figure creation were performed with GraphPad Prism 9.0.2. Each single measured/calculated value and mean ± SEM were plotted in the graphs unless otherwise specified. Statistical methods used were described in the Figure Captions. Statistical tests were two-tailed whenever appropriate.

## Supplementary Materials

Fig. S1. Glycomics analyses of human bronchial epithelial (HBE) and submucosal gland (SMG) mucus

Fig. S2. Sulfated glycan analyses of human bronchial epithelial (HBE) and submucosal gland (SMG) mucus

Fig. S3. PRR4 distribution in disease airways

Fig. S4. Region-specific localization of submucosal gland candidate biomarkers

Fig. S5. Absence of PRR4 degradation by co-incubation with human neutrophil elastase

Fig. S6. Validatory experiment results for anti-MUC5B antisera raised in-house

Data file S1. O-Glycans detected and quantified following release from HBE and SMG Mucin

Data file S2. Sulfated O-Glycans detected and quantified following release from HBE and SMG mucin

Table S1. Solubility of SMG mucus after addition of IGEPAL

## Acknowledgements

The authors thank the UNC CF Center Tissue Procurement and Cell Culture Core for providing human airway tissues and primary cells, and the UNC Animal Histology Core for paraffin embedding and sectioning the tissues. We also thank Dr. Hong Dang for statistical analyses and Eric Roe for editing the manuscript.

## Author contributions

T.K., S.T.B., B.B. and R.C.B. conceived the project and designed experiments. S.H.R. provided human airway tissues and primary cells. T.K., G.V.S. and S.T.B. collected pig SMG mucus. T.K. collected human SMG mucus. T.K., H.P.G, H.T., H.T.L and B.B. performed %solids, osmotic pressures, and cohesion measurements. G.V.S. performed solubility and light scattering assays. K.A., M.P. and M.T. performed glycomics analyses. M.J.P. and R.F. acquired SEM and immunofluorescent images of SMG mucus strands and characterized their structural properties. K.A.B. and K.K.W. acquired EM images of porcine and human airway tissues. K.O. prepared human histology sections and RNA samples from human airway tissues. R.C.G performed RNA *in situ* hybridization. T.K., S.M.B.C. and T.A.B. performed immunostaining and RT-qPCR. G.R. and M.K. performed proteomic analysis. M.R.M, C.B.M. and D.B.H performed analyses of mucus flakes. C.B.M and C.E. provided mouse anti-MUC5B antiserum. L.J.F. performed PRR4 overexpression and purification. T.N. and M.R. provided feedback to overall work. T.K., B.B., W.K.O. and R.C.B. analyzed and interpreted the data. B.B. and R.C.B. supervised the project. T.K., R.F., W.K.O, B.B. and R.C.B drafted and finalized the manuscript.

## Data and materials availability

All data are available in the main text or the supplementary materials.

---

**Figure S1.**
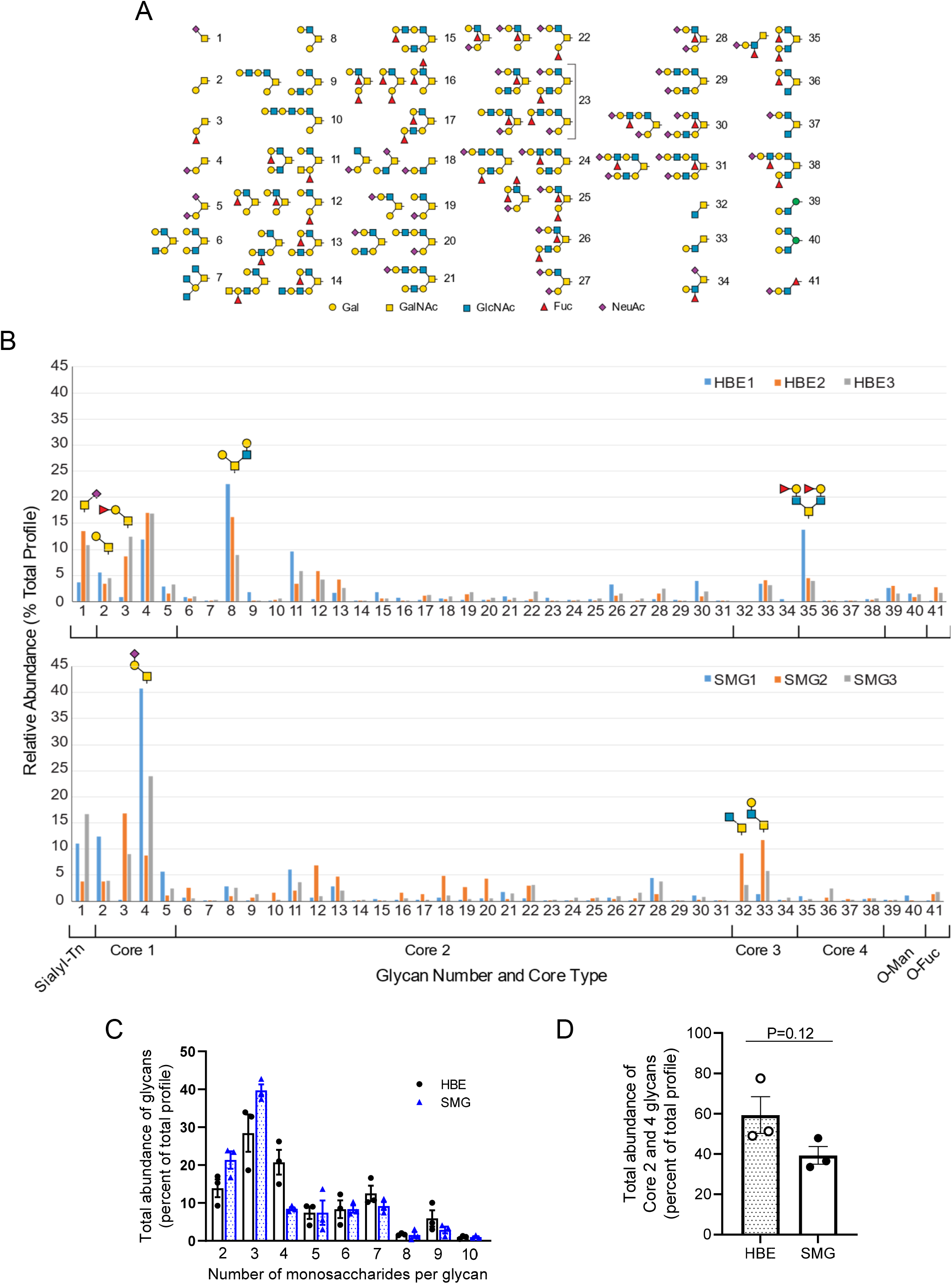
Glycomics analyses of human bronchial epithelial (HBE) and submucosal gland (SMG) mucus. **(A)** Glycan types detected in HBE and SMG mucus samples. **(B)** Relative abundance of each glycan type in HBE vs SMG mucus from the matched donors. **(C)** Abundance of glycans with 2 to 10 monosaccharides per glycan in HBE vs SMG mucus from matched donors. **(D)** Abundance of more complex (Core 2 and 4) in HBE vs SMG mucus. N = 3 for each. Unpaired t-test.

**Figure S2.**
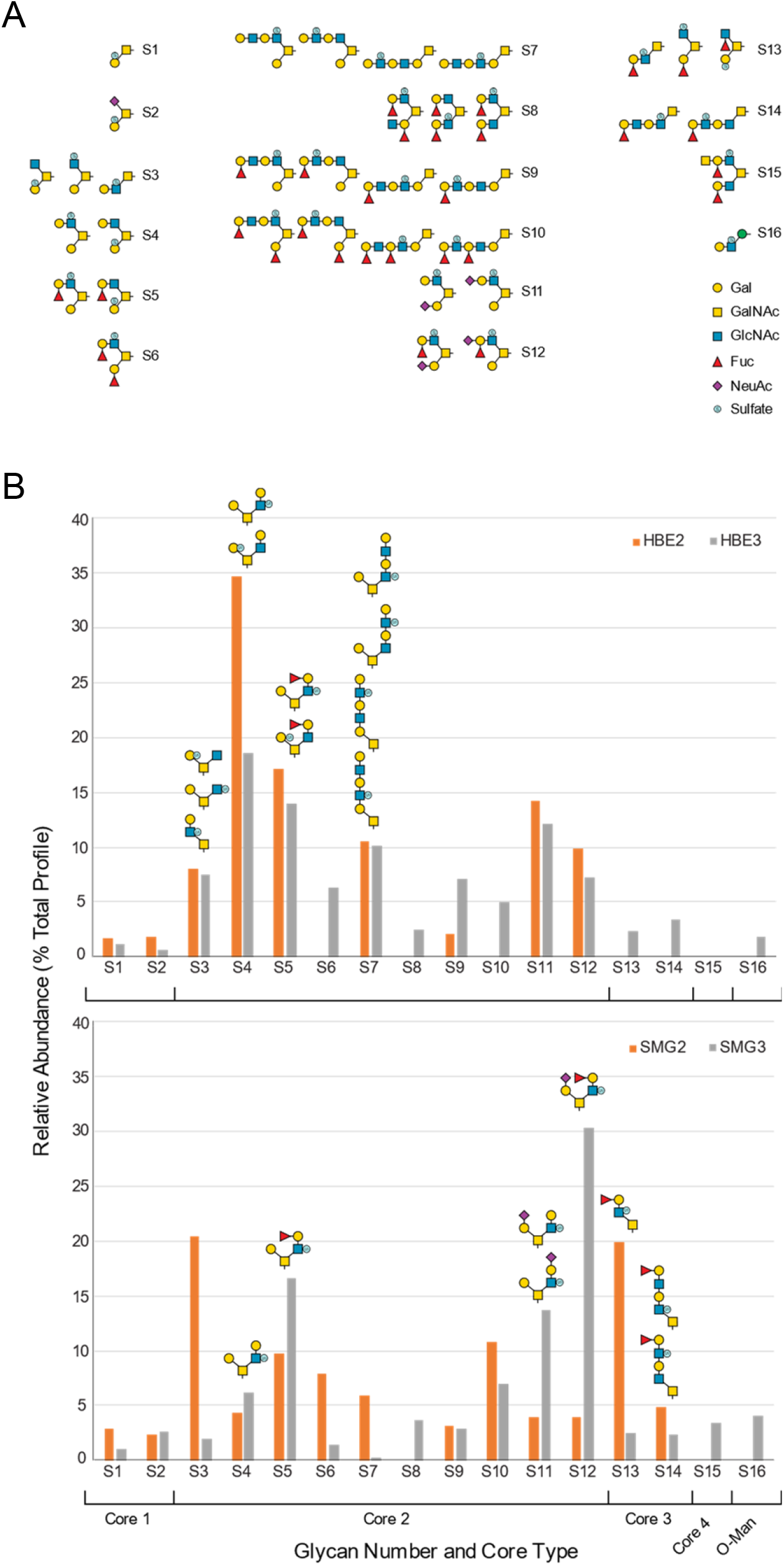
Sulfated glycan analyses of human bronchial epithelial (HBE) and submucosal gland (SMG) mucus. **(A)** Sulfated glycan types detected in HBE and SMG mucus samples. **(B)** Relative abundance of each sulfated glycan type in HBE vs SMG mucus from the matched donors. N = 2.

**Figure S3.**
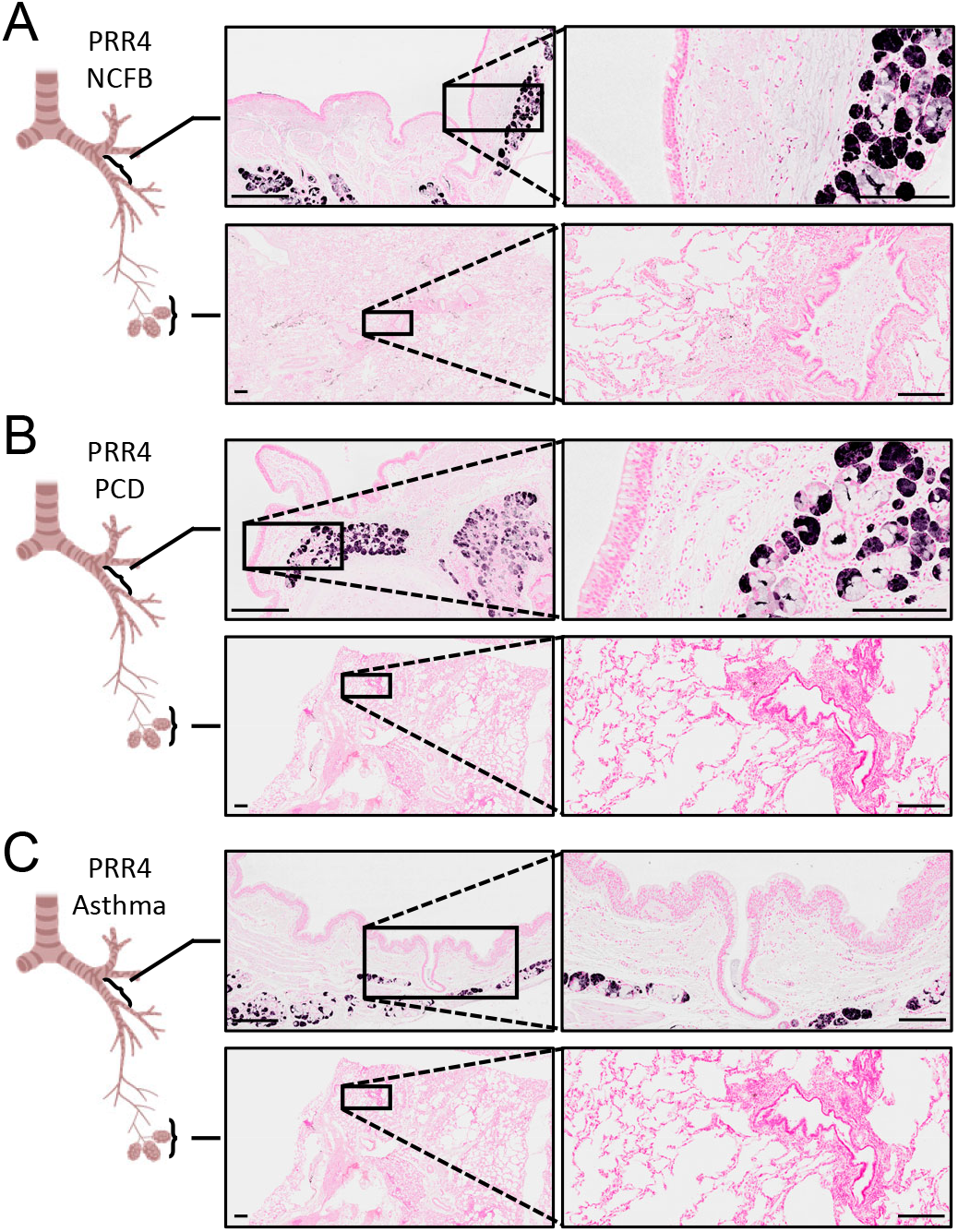
PRR4 distribution in disease airways. Immunohistochemistry for PRR4 in airways from (**A**) non-CF bronchiectasis (NCFB), (**B**) primary ciliary dyskinesia (PCD) and (**C**) asthma subjects. Bars: 500μm (low power images) and 200μm (magnified images).

**Figure S4.**
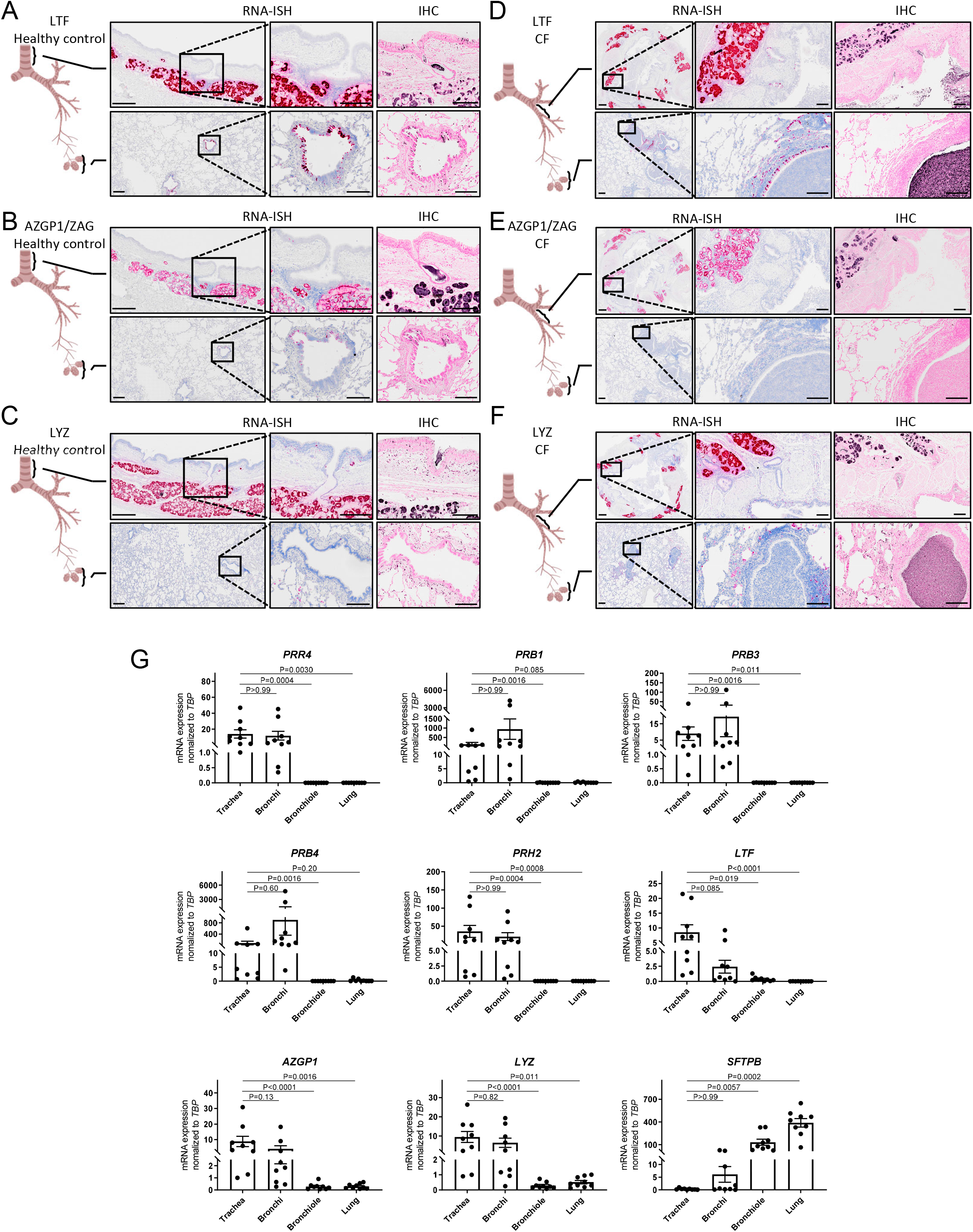
Region-specific localization of submucosal gland candidate biomarkers. (**A-F**) RNA *in situ* hybridization (RNA-ISH) and immunohistochemistry (IHC) for submucosal gland markers: (**A,D**) lactoferrin (LTF); (**B,E**) zinc-α2-glycoprotein (ZAGP1/ZAG); and (**C,F**) lysozyme (LYZ) in (**A-C**) normal and (**D-F**) CF human airways. Bars: 500μm (low power images) and 200μm (magnified images). (**G**) Quantitation of mRNA expression of candidate SMG marker genes including proline-rich protein family genes (*PRR4*, *PRB1*, *PRB2*, *PRB4* and *PRH2*), *LTF*, *AZGP1* and *LYZ*. A distal airway marker (*SFTPB*) was measured as a validation for airway region specificity. N = 9 donors. Friedman tests followed by Dunn’s multiple comparison tests.

**Figure S5.**
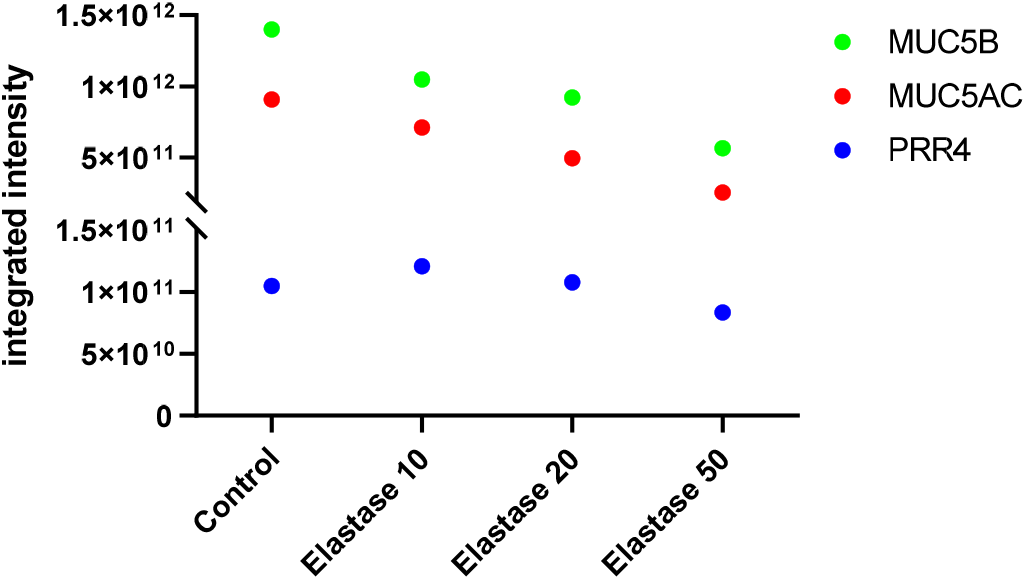
Absence of PRR4 degradation by co-incubation with human neutrophil elastase. Integrated intensity of MUC5B, MUC5AC and PRR4 peptides after incubation with human neutrophil elastase at designated concentrations for overnight as measured by mass spectroscopy, showing PRR4 resistance to HNE.

**Figure S6.**
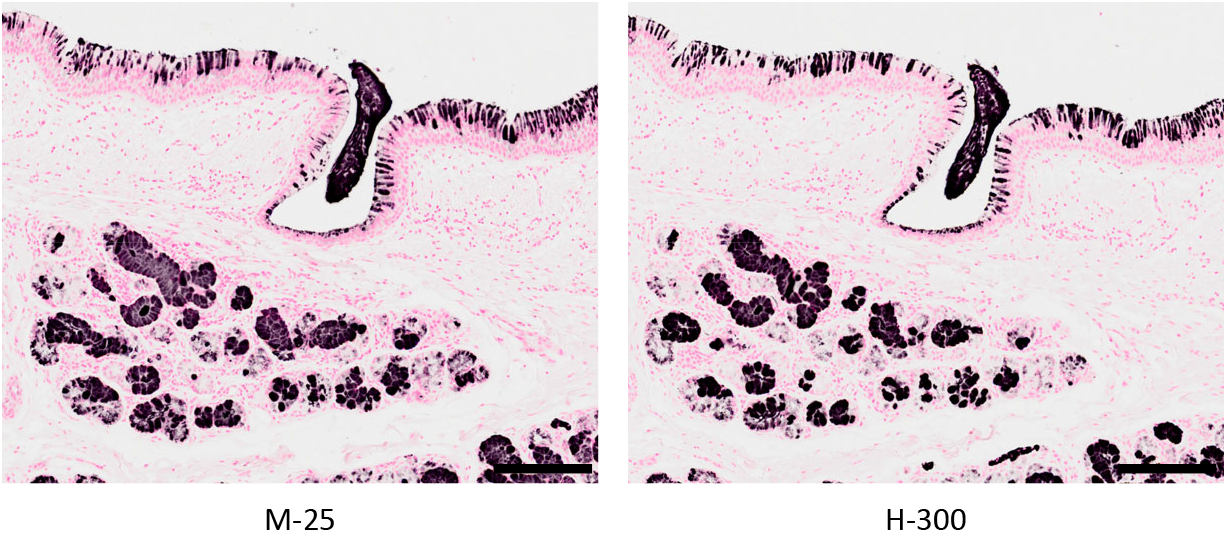
Validatory experiment results for anti-MUC5B antisera raised in-house. Immunohistochemistry for MUC5B with mouse antisera raised in-house (M-25) and rabbit polyclonal antibody (H-300) in healthy human trachea. Bar = 200μm.

**Supplementary Table 1. Solubility of SMG mucus after addition of IGEPAL**

All the PBS-insoluble components of human and pig SMG mucus exhibited full or partial solubility after adding IGEPAL.

